# Motor Imagery Improves Force Control in Older and Young Females

**DOI:** 10.1101/2024.10.02.616364

**Authors:** Cori A Calkins, Sarah N Kraeutner, Chris J McNeil, Jennifer M Jakobi

**Affiliations:** Healthy Exercise and Aging Lab, School of Health and Exercise Sciences, University of British Columbia Okanagan, Kelowna, British Columbia, Canada; Neuroplasticity, Imagery, and Motor Behaviour Lab, University of British Columbia, Kelowna, Canada; Djavad Mowafaghian Centre for Brain Health, University of British Columbia, Vancouver, Canada; Integrative Neuromuscular Physiology Lab, School of Health and Exercise Sciences, University of British Columbia Okanagan, Kelowna, British Columbia, Canada; Institute for Healthy Living and Chronic Disease Prevention, University of British Columbia Okanagan, Kelowna, British Columbia, Canada

**Author notes:** Corresponding Author: Jennifer (Jenn) Jakobi, PhD, Faculty of Health and Social Development Professor, Health and Exercise Sciences, The University of British Columbia – Okanagan campus 145-1147 Research Road (Arts Building), Kelowna, BC Canada V1V 1V7.

## Abstract

Motor imagery training (MIT) is the mental rehearsal of a motor task with no overt movement that enhances physical performance through adaptations in neural excitability. MIT may prime the motor system for physical execution. In older adults, with physical practice, force steadiness (FS) improves and changes are related to improved performance of functional tasks, and associated with adaptations in neural excitability. The purpose of this study was to determine if one session of MIT influences corticospinal excitability and improves FS of isometric elbow flexion contractions in young and older female adults. To test the hypotheses that MIT would increase corticospinal excitability and improve isometric elbow flexion FS to a greater extent in older compared to young females fourteen older (67-89 years old) and twenty-two younger (19-33 years old) participants were randomly assigned to a MIT group or Control group. Participants, in a block design, performed isometric elbow flexion contractions at 10% of maximal force prior to and following MIT (training group) or no training (Control group). Elbow flexion contractions were performed in blocks 1, 3, and 5. MIT or documentary viewing was performed in blocks 2 and 4. Motor evoked potentials (MEPs) elicited by transcranial magnetic stimulation were collected within the last five seconds of each submaximal contraction. The MEPs were reduced in the older MIT group from block 1 to block 5 (p=0.039) but not the young MIT group (p=0.761). Force steadiness in the older (p=0.005) and young (p=0.001) females improved from baseline after 20 minutes of MIT. Older females improved force steadiness relative to the baseline to a greater extent than young females (older 8.44% and young 5.0%), and the improvements were significant in older females in the first 10 minutes. In older females, MIT primes the motor system and improves FS earlier and to a greater extent than in young females.

## Introduction

The ability to maintain a steady contraction is essential for the execution of many motor behaviours. To estimate the stability of motor output, force steadiness (FS), quantified as the amplitude of force fluctuations around a target force during a submaximal isometric contraction, is often measured. FS of the elbow flexors is influenced by multiple factors, such as target force (Brown et al., 2010; Enoka et al., 2003), the position of the forearm (Brown et al., 2010; Smart, Kohn, et al., 2018), absolute strength (Brown et al., 2010; Pereira et al., 2015), and, notably, sex and age, as females are less steady than males (reviewed in Jakobi et al., 2018) and older adults are less steady than young (Feeney et al., 2018; Pereira et al., 2015; Smart, Baudry, et al., 2018; Tracy et al., 2007). Hence, older females are the least steady of these groups, potentially impeding their ability to perform functional tasks, as FS is related to functional ability (reviewed in Pethick et al., 2022). Therefore, it is important for older adults, especially females, to maintain or improve FS. The potential improvement of steady contractions in older females might minimize the exacerbated age-related decline in functional abilities reported for females compared with males (Butler et al., 2009).

One potential approach to improve FS is motor imagery training (MIT), which is the mental rehearsal of a motor task with no concurrent motor output (Collet et al., 2011). MIT priming prepares the motor system for motor task execution. It has been demonstrated that MIT improves force control in young adults (Fukumoto et al., 2021; Tatemoto et al., 2017) as well as increases neural drive (Grosprêtre et al., 2018) and corticospinal excitability (Kasai et al., 1997; Lebon et al., 2012; Lee et al., 2021; Mizuguchi et al., 2013). However, it is unknown if MIT leads to improvements in force steadiness and if there will be a concurrent increase in corticospinal excitability in older and young adults. Greater motor imagery ability is associated with greater corticospinal excitability during MIT (Lebon et al., 2012). Older adults have reduced motor imagery quality (Malouin et al., 2010) and motor imagery ability (Kalicinski et al., 2015; Muto et al., 2022) compared to young adults. However, if motor imagery ability is sufficient in older females, it might offer a means to enhance FS through alterations in corticospinal excitability.

If MIT priming occurs and the produced effects are favourable, MIT could have a meaningful influence on the performance of steady movements in young females as well as in older females who experience declines in FS with age. The primary aim of this study was to determine if one session of MIT produces increases in corticospinal excitability and improvement to FS of isometric elbow flexion contractions in young and older females. The secondary aim was to ascertain how motor imagery ability and potential differences between young and older females contribute to changes in FS. It was hypothesized that MIT will increase corticospinal excitability and improve FS, with the greatest benefit observed in older females.

## Methods

### Participants

Thirty-eight females aged 19-35 and 65-90 years of age volunteered for this study. One young participant withdrew between familiarization and the experimental day. One older participant was excluded due to the inability to perform the force-tracking task. Fourteen older (n=7 Control, n=7 MIT) and twenty-two young participants (n=11 Control, n=11 MIT) completed the study. All participants were self-reported right-hand dominant, with no known neurological disorders. Participants were excluded if they had a self-reported history of MIT or training in fine motor tasks (e.g. musicians), high levels of resistance training, or had injury or surgery to the right arm in the prior 6 months. Individuals with severe cognitive impairment, unable to read or speak English fluently, or with any contraindication to TMS (Rossi et al., 2009) were also excluded. Before participation in the study, all participants gave informed written consent to a protocol approved by the institution’s Clinical Research Ethics Board (H22-01542).

Participants were randomly assigned to the MIT or Control group. Participants visited the laboratory on two occasions: familiarization day and experimental day (described below). To account for hormonal fluctuations on the experimental day, testing was during periods when estrogen and progesterone were at their lowest. That is, young females not using hormonal contraceptives, as well as those using oral contraceptives, were tested between days 1-7 of the menstrual cycle. One young participant was not tested during these days (menses) due to wildfires in Kelowna BC. If using an IUD, participants were tested at any time point. Older post-menopausal females were tested at any time period, and none reported hormonal replacement therapy.

### Experimental Setup and Procedures

Participants were seated in a custom dynamometer chair that was adjusted to maintain hip and knee flexion at 90 degrees, with the right arm abducted to 15 degrees and 100 degrees of flexion at the elbow. Participants were secured to the chair with a seatbelt-like restraint for consistent body positioning and isolation of elbow flexion contractions. Participants grasped the manipulandum with their right hand, with their forearm and hand in a neutral position (thumb up). The force transducer was positioned underneath the manipulandum to measure elbow flexion force. To ensure visual feedback was consistent, force signals were displayed on a 52-cm computer monitor located 100cm in front of the participant, at eye-level. EMG signals were hardware amplified by 100 (Coulbourn Instruments, Massachusetts, USA). EMG signals were sampled at 2008Hz and force signals were sampled at 1004Hz using a 1401plus analog-to-digital converter (Cambridge Electronic Design, Cambridge, UK) and stored for offline analysis using Spike 2 software (version 10; CED, Cambridge, UK).

### Familiarization Day

Following informed consent and screening, participants completed the Paffenbarger Physical Activity Questionnaire (Paffenbarger et al., 1993). Participants in the MIT group also completed the Movement Imagery Questionnaire-Revised Second Edition (MIQ-RS) (Gregg et al., 2010) to characterize an individual’s motor imagery ability on familiarization day. The MIQ-RS has two 7-point Likert scales: visual (“see”) and kinesthetic (“feel”); Participants are asked to self-report using these scales to assess how difficult or easy it was to mentally rehearse a visual or kinesthetic item from the MIQ-RS; One is difficult to see or feel and seven is easy to see or feel. Unlike items from the MIQ-RS, this study’s mental rehearsal task involved both visual and kinesthetic motor imagery concurrently. Therefore, to characterize an individual’s motor imagery ability during this study’s experimental day, participants used both of these self-report scales after each mental rehearsal time period. We also obtained mental chronometry for each item on the MIQ-RS. Participants self-timed both the physical tasks and the duration of their motor imagery using a stopwatch.

Voluntary activation was established with the interpolated twitch technique (Allen et al., 1995) using muscle belly stimulation of the biceps brachii by applying transcutaneous electrical stimulation (DS7AH, Digitimer Ltd., Welwyn Garden City, UK) through carbon rubber stimulation electrodes (4 × 4.5 cm) placed over the muscle belly. While participants were at rest, the current intensity of single, square-wave stimulations (200μs pulse width, ≤400V) was increased until no further increase in elbow flexion twitch force was observed, and then increased a further 15% from this level to achieve supramaximal stimulation intensity. A maximal voluntary contraction (MVC) without stimulation was undertaken for familiarization, and then two to three MVCs were executed, with three single-pulse stimulations (200μs pulse width) applied before, during, and after the MVC. Approximately two minutes of rest separated the MVCs to prevent muscle fatigue. All participants performed a minimum of three MVCs and then practiced three isometric elbow flexion tracking tasks at 10% MVC.

During the FS tasks, participants in the Control and MIT group were provided a target line and instructed through an audio speaker when to begin and end each force task. The audio recording had an audible ‘up’ to increase force during the ramp to target (∼ two seconds), a clear ‘hold’ for when to plateau and maintain the target force for 25 seconds, and then the ‘rest’ command was given to de-ramp (∼two seconds) and relax. Each FS task was separated by 10 seconds of rest. These instructions were verbally explained before commencing the protocol. The MIT group was given a motor imagery familiarization script (see Supplementary Appendix A) to read and familiarize themselves with MIT. The group then watched a short video from the first-person perspective (i.e., over the shoulder) of the 10% MVC force task that depicted the right (dominant) limb and force output on a computer monitor. Participants in the MIT group were then asked to relax and practice mentally rehearsing three times the 10% MVC force task using both kinesthetic and visual imagery adopting a 1^st^ perspective to start to form a picture in their head of this study’s force task. Participants did not have any audio instruction for when to begin and end each mental rehearsal of the task.

### Experimental Day

Surface electrodes (4mm, Ag-AgCl) were positioned over the biceps brachii muscle belly and the distal biceps brachii tendon, with the ground electrode placed on the bony prominence of the olecranon and scapula. To elicit a compound muscle action potential (M-wave) from the biceps brachii, one electrode was placed over the brachial plexus at Erb’s point for attachment of the cathode and one over the acromion process for attachment of the anode. With the forearm in a neutral position, stimulation intensity of the DS7AH was established by applying single, square-wave pulses (200μs pulse width) to the brachial plexus, and incrementally increasing the current until no further increase of M-wave peak-to-peak amplitude was found (Mmax). Stimulation intensity was then multiplied by 1.15 to establish the supramaximal intensity. Three stimulations at supramaximal intensity were conducted approximately 10 seconds apart, and, to determine 10% Mmax, the average of the Mmax peak-to-peak amplitude was multiplied by 0.1.

During all TMS procedures, participants wore a cloth TMS cap (Caputron, New Jersey, USA). The vertex was located, and marked as position 1 on the cap, by measuring the midpoint between the nasion and inion as well as the midpoint between the tragi. One centimeter was measured anterior, anterior-lateral, lateral, posterior-lateral, and posterior to the vertex, with these positions marked as 2 to 6 on the cap. As all participants were right-handed, lateral shifts were to the left of the vertex; the coil was oriented to stimulate the left hemisphere. With the use of a round coil (90mm) and a Magstim 200^2^ (Magstim Ltd., Wales, UK), single-pulse TMS was delivered over positions 1-6 to stimulate the primary motor cortex. The optimal stimulation site was established by setting the TMS intensity to 60% and determining which position had the greatest MEP peak-to-peak amplitude from biceps brachii while participants were at rest. Once the optimal stimulation site was located, the rim of the TMS round coil was marked on the cap so placement of the coil would remain consistent throughout testing. TMS intensity was determined by having participants contract at 10% MVC and then incrementally adjusting TMS intensity until MEP peak-to-peak amplitude was ∼10% Mmax.

The experimental day was split into five blocks. The Control group and MIT group were tested in a matched-block design, and FS was measured for an identical number of contractions. Participants in the Control group and MIT group first performed two to three MVCs before the testing blocks (Figure 1). In blocks 1, 3, and 5, all participants performed seven FS tracking tasks at 10% MVC with audio instructions, as described above on familiarization day. Within the last 5 seconds of the steady-state phase of each contraction, a variable single-pulse TMS was delivered to the stimulation site. In blocks 2 and 4, the MIT group was instructed through a speaker to mentally rehearse the isometric tracking tasks at 10% MVC without any overt movements. Participants were instructed to visually and kinesthetically mentally rehearse the task from a first-person perspective from a relaxed seated position in the chair with their dominant arm relaxing in the elbow stand and hand not firmly gripping the manipulandum. Participants were monitored by the researchers during MIT to ensure no overt movements were performed during mental rehearsal. MIT was conducted in blocks consisting of 10 minutes of mental rehearsal with 2 minutes of rest in between to prevent mental fatigue. As part of the mental rehearsal time, participants listened to an MIT script (see Supplementary Appendix B) at the beginning and after the rest period of each MIT block. Each MIT block had a total of 14 trials that involved 25 seconds of mental rehearsal at the 10% MVC target force followed by 10 seconds of rest. For each trial of MIT, participants were signaled with the same audible ‘up’ for ramp to target, ‘hold’ for when to plateau at the target force, and ‘rest’ for when to de-ramp and relax as the force tasks for each trial of MIT. At the 5- and 10-minute time points of blocks 2 and 4, participants rated themselves visually and kinesthetically with the 7-point Likert scale. The Control group was time-matched to MIT and watched The Green Planet: Water Worlds for 10 minutes with 2 minutes of rest in between for a total of 12 minutes for each of blocks 2 and 4.

**Figure 1.**
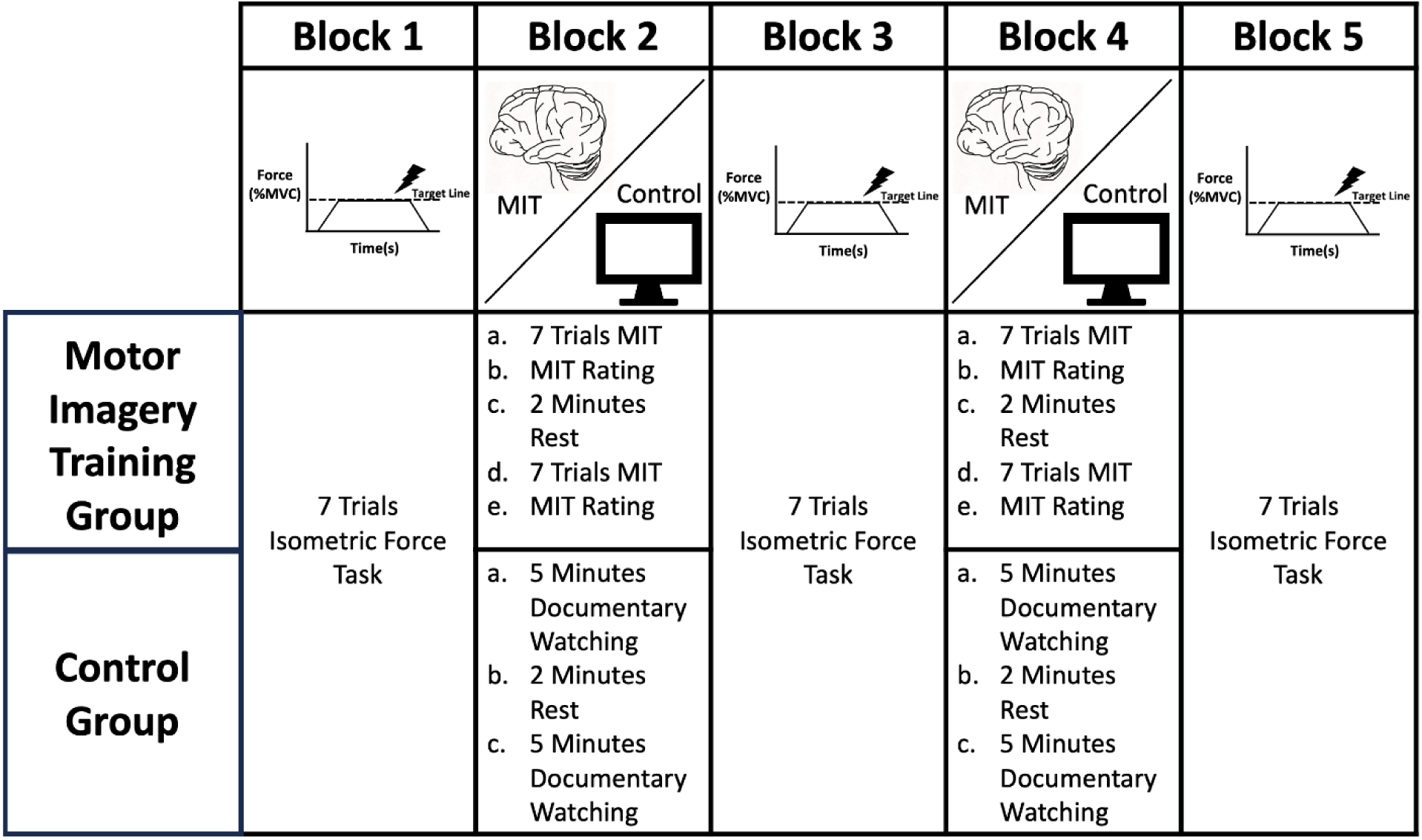
Overview of Experimental Day Block Order. Blocks 1, 3, and 5 involved the 10% MVC force tracking tasks for the MIT group and Control group. Blocks 2 and 4 involved MIT or watching a documentary for 10 minutes. At the 5-minute and 10-minute time points of blocks 2 and 4, the MIT group participants rated their visual and kinesthetic motor imagery ability. MIT, motor imagery training; %MVC, percentage of maximal voluntary contraction

Thus, the series of blocks were ordered: 1) isometric elbow flexion tracking tasks, 2) MIT or documentary (i.e., The Green Planet: Water Worlds) viewing (MIT vs. Control group), 3) isometric elbow flexion tracking tasks, 4) MIT or documentary viewing, 5) isometric elbow flexion tracking tasks. All blocks were of identical duration (Figure 1).

### Data Analysis

From the twitch interpolation technique, voluntary activation for each participant was calculated as: VA= [1 − (Interpolated Twitch Force ÷ Resting Twitch Force)] × 100%. Using a custom MATLAB script, force steadiness was analyzed as the coefficient of variation (CV) of the force around the target force level during the 20-second plateau before the TMS pulse of each tracking task. CV of force was calculated with the equation CV (%) = (Standard Deviation of Force ÷ Mean of Force) × 100%. Spike 2 version 10 (CED, Cambridge, UK) was used to analyze evoked force and EMG. Peak-to-peak MEP amplitude was measured for the biceps brachii during each isometric elbow flexion tracking task at 10% MVC force.

Visual and kinesthetic imagery ability scores from the MIQ-RS were calculated as the average of 7 items for each participant. Global imagery ability scores from the MIQ-RS were calculated from the average of the 7 visual and 7 kinesthetic items for each participant. Mental chronometry was assessed as the time difference between the action time and mental task time for each item on the MIQ-RS. Average visual, kinesthetic, and global mental chronometry were calculated for each participant. Kinesthetic, visual, and global motor imagery ability scores after 5 minutes and 10 minutes of each MIT block (i.e., blocks, 2 and 4), were summed for a total block score, as well as averaged to characterize each block separately and overall training.

### Statistical Analyses

All correlation, t-test, and ANOVA analyses were conducted using the software SPSS version 29 (IBM, Armonk, New York, USA), whereas all linear mixed effects (LME) models were conducted using the software R (R version 4.3.2, The R Foundation for Statistical Computing). Mauchly’s test of sphericity was undertaken for each ANOVA, and a Greenhouse-Geisser correction was used if p<0.05. For FS (CV of force), six trials (6/292; 2%) from the older and 11 trials from the young adults (11/459; 2%) were removed and for MEPs, one trial from the older (1/263;0.4%) and three trials from the young adults (3/452;0.7%) were removed for being 3x greater than interquartile range. In the ANOVA and correlation analyses extreme outliers were replaced with winsorized (1.95 × SD + mean) data, and this occurred for the MIQ-RS mental chronometry for one older participant, in the MIT group.

#### Age Effects

To test for baseline group differences, a 2 (Age; young, older) × 2 (Group: MIT, Control) two-way ANOVA was conducted to compare participant characteristics of anthropometrics, strength, voluntary activation, physical activity, Mmax amplitude and CV of Force at baseline (Block 1). A one-tailed independent-sample t-test was conducted to compare young and older female’s mean MIQ-RS mental chronometry from visual, kinesthetic, and global items. The relationship between motor imagery ability relative to the time difference between performance and mental rehearsal of a task in the young and older females was also evaluated with Pearson r correlational analyses between the self-rated MIQ-RS scores (visual, kinesthetic, and global) and mental chronometry (visual, kinesthetic, and global).

#### Motor Imagery Training Effects

To address the first hypothesis that MIT will increase FS and corticospinal excitability as well as the second hypothesis that the change would be greater in older females separate linear mixed effects (LME) models were conducted for older and young adults for CV of Force and MEPs. The LME models included group, block, and the interaction of group and block as fixed effects, with the participant as a random intercept. Group was sum-contrast coded, and the block was treatment-coded to make the required comparisons for our hypotheses. LME models with Satterthwaite’s method were fit with packages lme4 (Bates et al., 2015) and lmerTest (Kuznetsova et al., 2017). Cohen’s d values between blocks were computed with the package effsize (Torchiano, 2020).

To understand why older females would improve to a greater extent than young females, motor imagery ability between older and young females over the training session, block 2 and 4 were compared with a 2 (Age) × 2 Block (block 2, block 4) mixed-factorial ANOVA for kinesthetic, visual and global MIT scores. The influence of the blocks of MIT on CV of force and corticospinal excitability was explored by calculating the percent change in CV of force and MEPs between block 1 and block 3 as well as block 1 and 5, then using a Person r correlation analysis between the independent variables of MIT (kinesthetic, visual and global) and the change in CV of force.

## Results

### Age Effects

Fourteen older (n=7 Control, n=7 MIT) and 22 young female participants (n=11 Control, n=11 MIT) completed the study (Table 1). The young were stronger than the older females (p=0.002), had higher physical activity scores (p=0.003), and were steadier at baseline (p=0.03). The young did not differ from older females in voluntary activation scores (p=0.11) and Mmax amplitude (p=0.52). The young and older females in the MIT group did not differ from the Control group in strength (p=0.42), voluntary activation (p=0.54), physical activity scores (p=0.76), CV of force at baseline (p=0.09) and Mmax amplitude (p=0.92).

**Table 1.**
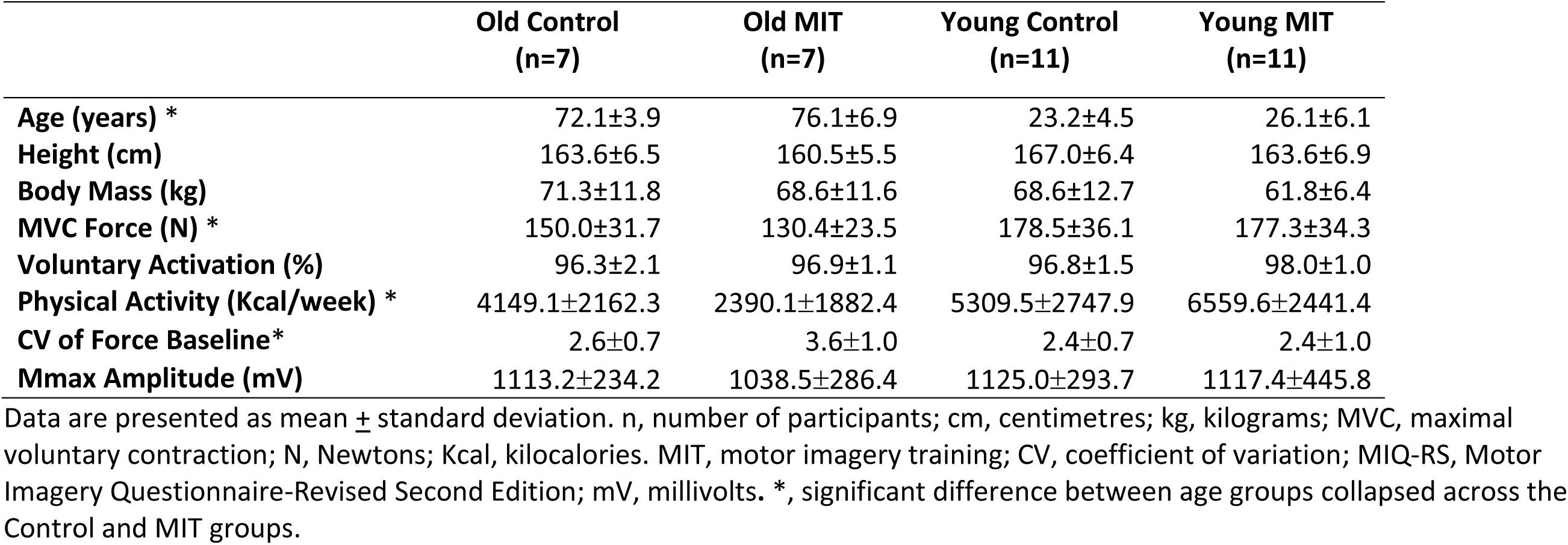
Participant Characteristics.

There was no significant block effect for kinesthetic, visual, or global MIT scores (p>0.05) or interactions for Age × Block for MIT scores (p>0.05) (Table 2). A distribution of MIQ-RS scores are shown in figure 2(a,b,c). No age differences were observed between kinesthetic (p=0.15), visual (p=0.39), and global (p=0.30) mental chronometry obtained from the MIQ-RS (Figure 2d). A main effect was observed for age for visual MIT scores (F(1,16)=4.456, ηp^2^=0.218, p=0.05) and global MIT scores (F(1,16)=5.26, ηp^2^=0.247, p=0.036), as young had higher values compared to older (Figure 3). A positive relationship was observed for global MIQ-RS scores and MIQ-RS global mental chronometry (r=0.756, p=0.007), but only in younger adults. Further, this positive relationship in the young adults was also observed for the kinesthetic scores (r=0.839, p=0.001). No other relationships between MIQ-RS scores and MIQ-RS mental chronometry were observed (p≥0.151).

**Figure 2.**
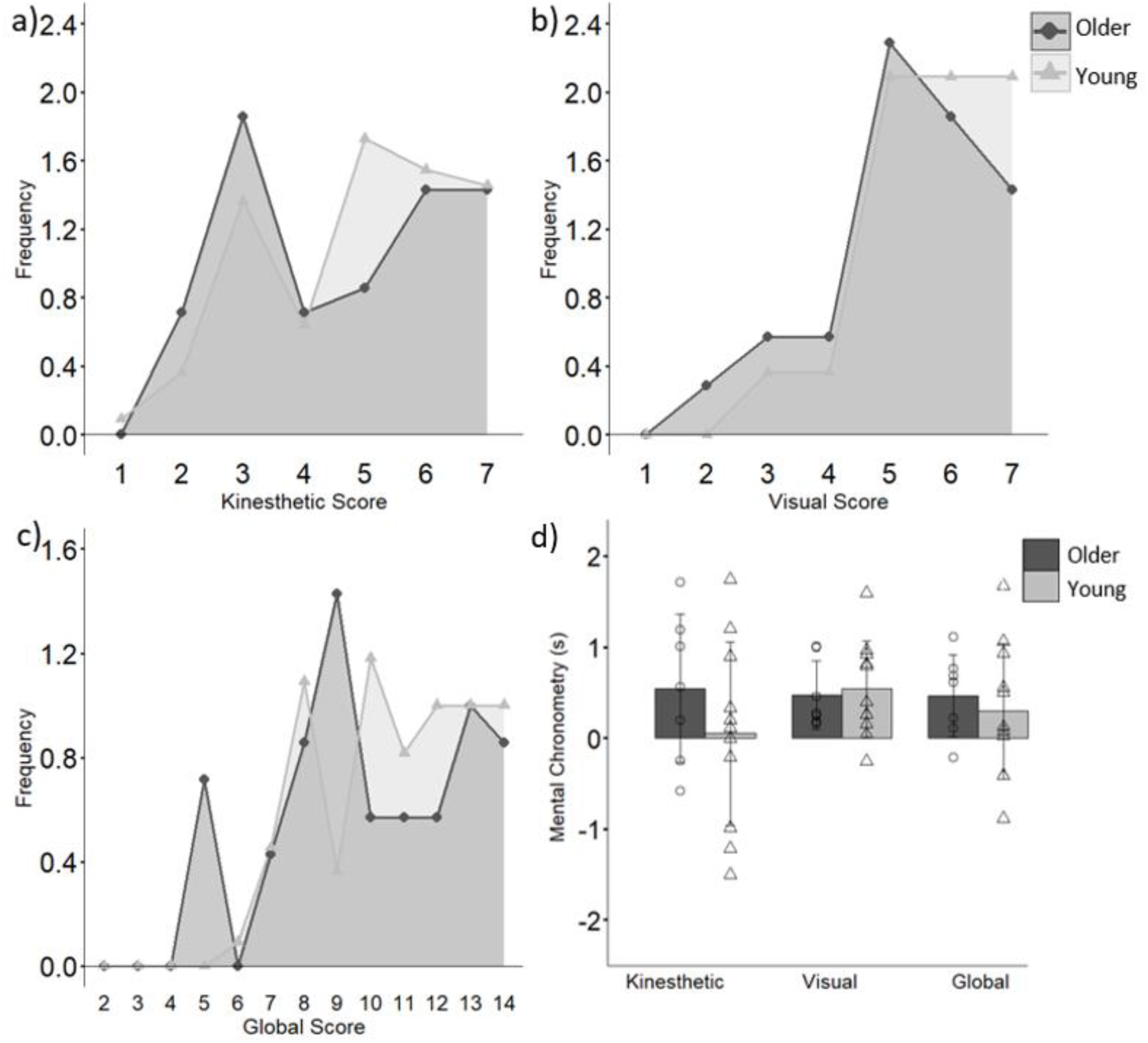
Motor imagery ability from the MIQ-RS for kinesthetic (a), visual (b) and global (c) imagery for older and young females. Frequency is the count from the MIQ-RS scores divided by the number of participants for each group. Mental chronometry (d) calculated as Action task time – Mental task time from the MIQ-RS items for older and young females. MIQ-RS, Motor Imagery Questionnaire-Revised Second Edition.

**Figure 3.**
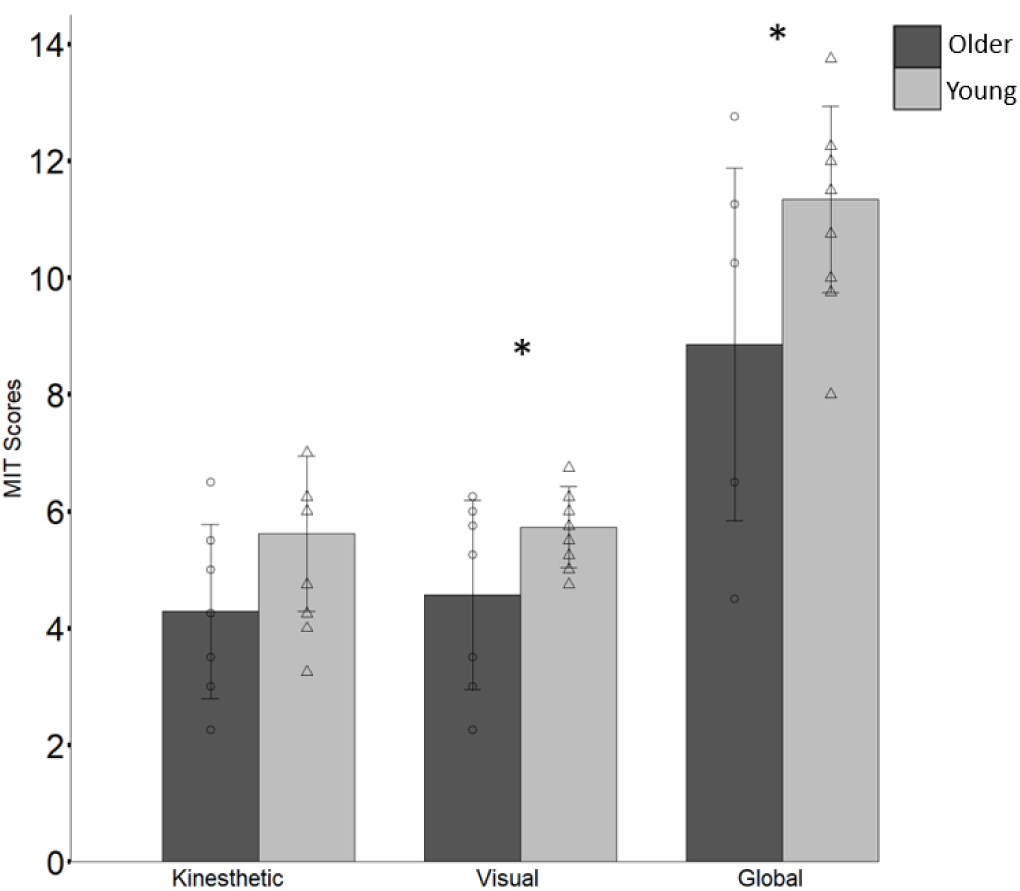
Motor imagery ability for young and older females. Mean kinesthetic, visual, and global MIT Scores from blocks 2 and 4. *, significant difference between young and older females. MIT, motor imagery training.

**Table 2.** MIT Kinesthetic, Visual, and Global Imagery Scores from blocks 2 and 4.

| Variables | Older (n=7)<br>Mean±SD | Young (n=11)<br>Mean±SD |
| --- | --- | --- |
| **Kinesthetic MIT Score/7** Block 2 | 4.43±1.10 | 5.55±1.51 |
| **Kinesthetic MIT Score/7** Block 4 | 4.14±2.10 | 5.68±1.25 |
| **Visual MIT Score /7** Block 2 | 4.64±1.46 | 5.77±0.68 |
| **Visual MIT Score/7** Block 4 | 4.5±1.94 | 5.68±0.81 |
| **Global MIT Score /14** Block 2 | 9.07±2.41 | 11.32±1.74 |
| **Global MIT Score /14** Block 4 | 8.64±3.93 | 11.36±1.57 |

Data are presented as mean± standard deviation. MIT, motor imagery training.

### MIT Training Effects

There was a significant improvement in the CV of force in the older MIT group from block 1 to block 3 (p=0.035), as well as block 1 to 5 (p=0.005). In the young MIT group, an improvement in CV of force was observed from block 1 to block 5 (p=0.001) (Table 3; Figure 4 a,b). Within the MIT group, the older had greater improvement in CV of force compared to the young between blocks 1 to 3 (d_older_= 0.33, d_young_= 0.07) and between blocks 1 to 5 (d_older_= 0.29, d_young_= 0.21). The peak-to-peak amplitude of MEPs was reduced in the older MIT group from block 1 to block 5 (p=0.039) (Table 3; Figure 4c). Within the MIT group, the young had a greater reduction in corticospinal excitability compared to the older between blocks 1 and 3 (d_older_= 6.34e^-04^, d_young_= 0.152). Whereas, the older MIT group had a greater reduction in corticospinal excitability compared to the young MIT group between blocks 1 to 5 (d_older_= 0.253, d_young_= 0.210 (Table 4; Figure 4 c, d).

**Figure 4.**
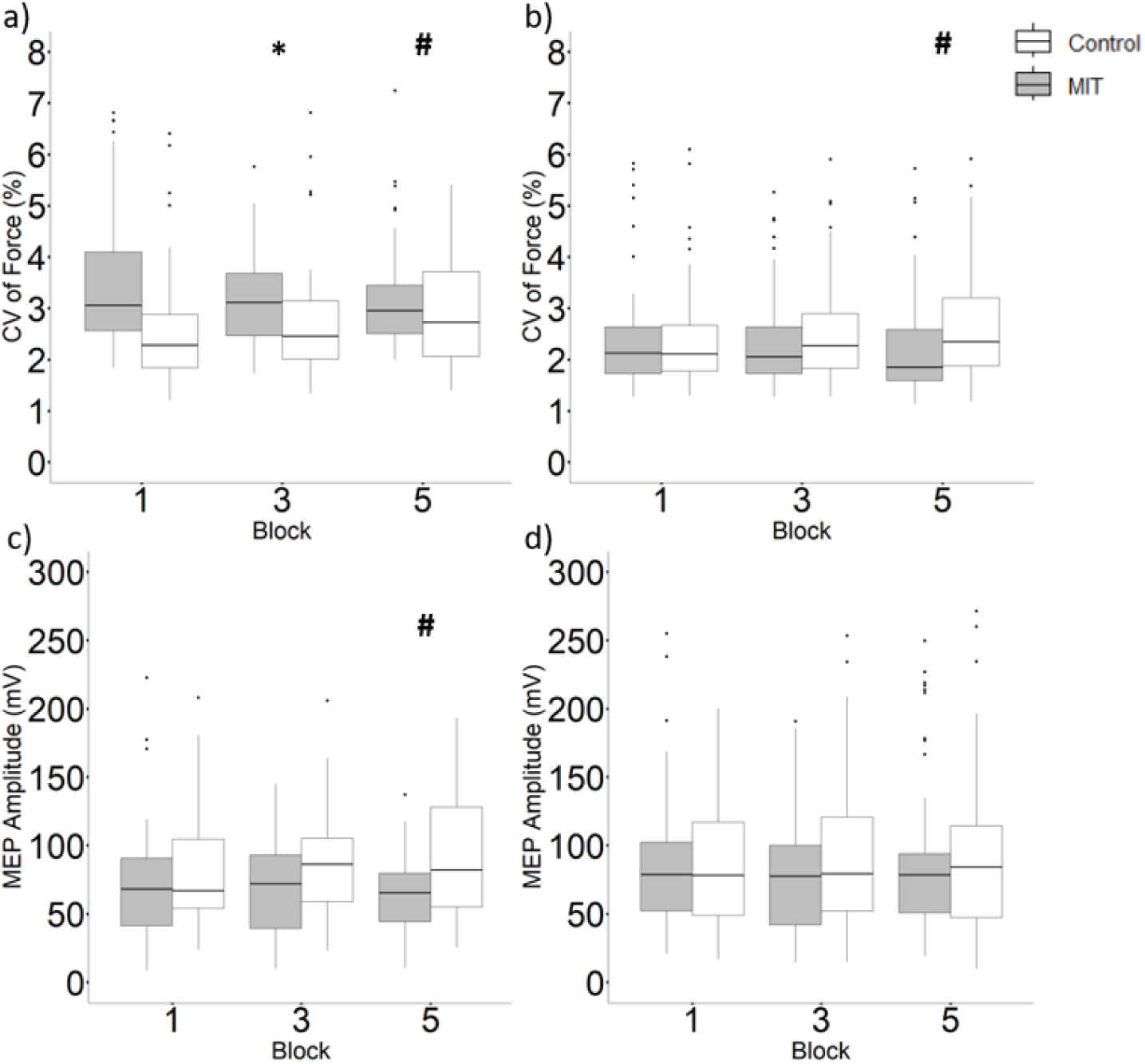
CV of force (a,b) and MEP amplitude (c,d) for older adults (a,c) and young adults (b,d) in the motor imagery training and Control groups for the 3 blocks that the submaximal tracking tasks occurred to quantify force steadiness. *Significant decrease between block 1 to 3; and # significant decrease between block 1 to 5. CV, coefficient of variation; MIT, motor imagery training; mV, millivolts volts.

**Table 3.** LME Model to Predict CV of Force in Older and Young Adults in MIT and Control Groups.

| Predictors | Older Adults |  |  | Young Adults |  |  |
| --- | --- | --- | --- | --- | --- | --- |
|  | Estimates | Standard Error | p | Estimates | Standard Error | p |
| **Fixed Effects** |  |  |  |  |  |  |
| (Intercept) | 3.085 | 0.213 | **1.470x10<sup>-10</sup>** | 2.398 | 0.198 | **5.370x10<sup>-11</sup>** |
| MIT | 0.509 | 0.213 | **0.029** | 0.007 | 0.198 | 0.972 |
| Block3 | -0.135 | 0.132 | 0.308 | 0.098 | 0.062 | 0.112 |
| Block5 | 9.787x10<sup>-4</sup> | 0.133 | 0.994 | 0.085 | 0.062 | 0.171 |
| MIT × Block3 | -0.281 | 0.132 | **0.035** | -0.089 | 0.062 | 0.152 |
| MIT × Block5 | -0.372 | 0.133 | **0.005** | -0.211 | 0.062 | **0.001** |
| **Random Effects** |  |  |  |  |  |  |
| Groups | Name | Variance | Standard Deviation | Name | Variance | Standard Deviation |
| Participants | (Intercept) | 0.510 | 0.714 | (Intercept) | 0.8209 | 0.9060 |
| Residual |  | 0.836 | 0.914 |  | 0.2842 | 0.5331 |
| Number of Observations | 286 |  |  | 448 |  |  |
| Number of Participants | 14 |  |  | 22 |  |  |

**Note: bolded text under the p column indicates significant (p <0.05) effects. MIT, motor imagery training**

**Table 4.** LME Model to Predict Corticospinal Excitability in Older and Young Adults in MIT and Control Groups.

| Predictors | Older Adults |  |  | Young Adults |  |  |
| --- | --- | --- | --- | --- | --- | --- |
|  | Estimates | Standard Error | p | Estimates | Standard Error | p |
| **Fixed Effects** |  |  |  |  |  |  |
| (Intercept) | 7.766x10<sup>-2</sup> | 9.975x10<sup>-3</sup> | **4.620x10<sup>-6</sup>** | 8.533x10<sup>-2</sup> | 9.254x10<sup>-3</sup> | **5.730x10<sup>-9</sup>** |
| MIT | -7.595x10<sup>-3</sup> | 9.975x10<sup>-3</sup> | 0.461 | -2.096x10<sup>-3</sup> | 9.254x10<sup>-3</sup> | 0.823 |
| Block3 | 2.074x10<sup>-3</sup> | 3.828x10<sup>-3</sup> | 0.589 | -7.973x10<sup>-4</sup> | 3.280x10<sup>-3</sup> | 0.808 |
| Block5 | -1.412x10<sup>-3</sup> | 3.794x10<sup>-3</sup> | 0.710 | 1.359x10<sup>-3</sup> | 3.273x10<sup>-3</sup> | 0.678 |
| MIT × Block3 | -8.993x10<sup>-4</sup> | 3.828x10<sup>-3</sup> | 0.815 | -4.725x10<sup>-3</sup> | 3.280x10<sup>-3</sup> | 0.150 |
| MIT × Block5 | -7.870x10<sup>-3</sup> | 3.794x10<sup>-3</sup> | **0.039** | -9.952x10<sup>-4</sup> | 3.273x10<sup>-3</sup> | 0.761 |
| **Random Effects** |  |  |  |  |  |  |
| Groups | Name | Variance | Standard Deviation | Name | Variance | Standard Deviation |
| Participants | (Intercept) | 0.001 | 0.035 | (Intercept) | 0.002 | 0.042 |
| Residual |  | 0.001 | 0.025 |  | 0.001 | 0.028 |
| Number of Observations | 263 |  |  | 452 |  |  |
| Number of Participants | 13 |  |  | 22 |  |  |

**Note: bolded text under p column indicates significant (p <0.05) effects. MIT, motor imagery training;**

A negative relationship was observed between global motor imagery scores of block 2 and CV of force block 3 in the older MIT group (Figure 5e; r=−0.678, p=0.047), yet not in the young MIT group (r=−0.185, p=0.293). There were no relationships observed between global motor imagery scores in block 4 and CV of force block 5 for older (r=−0.309, p=0.250) and young (r=−0.104 p=0.380) females (Figure 5f). There were no relationships evident between blocks for kinesthetic motor imagery scores in older (block 2 and 3 r=−0.569, p=0.091; block 4 and 5 r=−0.362, p=0.212) and young (block 2 and 3 r=−0.216, p=0.261; block 4 and block 5 r=−0.108, p=0.376) females with CV of force (Figure 5a and 5b). A negative relationship was observed between visual motor imagery scores of block 2 with CV of force block 3 in the old MIT group (Figure 5c; r=−0.6882^2^, p=0.044,), and no relationship for block 4 and CV of force block 5 (Figure 5d; r=−0.236, p=0.306). There were no relationships between visual motor imagery scores and CV of force in young (block 2, r=0.007, p=0.492; block 4, r=−0.035 p=0.459) (Figure 5c and 5d). There was no relationship between MEPs and the global, kinesthetic, and visual motor imagery scores across all block comparisons in older and young adults.

**Figure 5.**
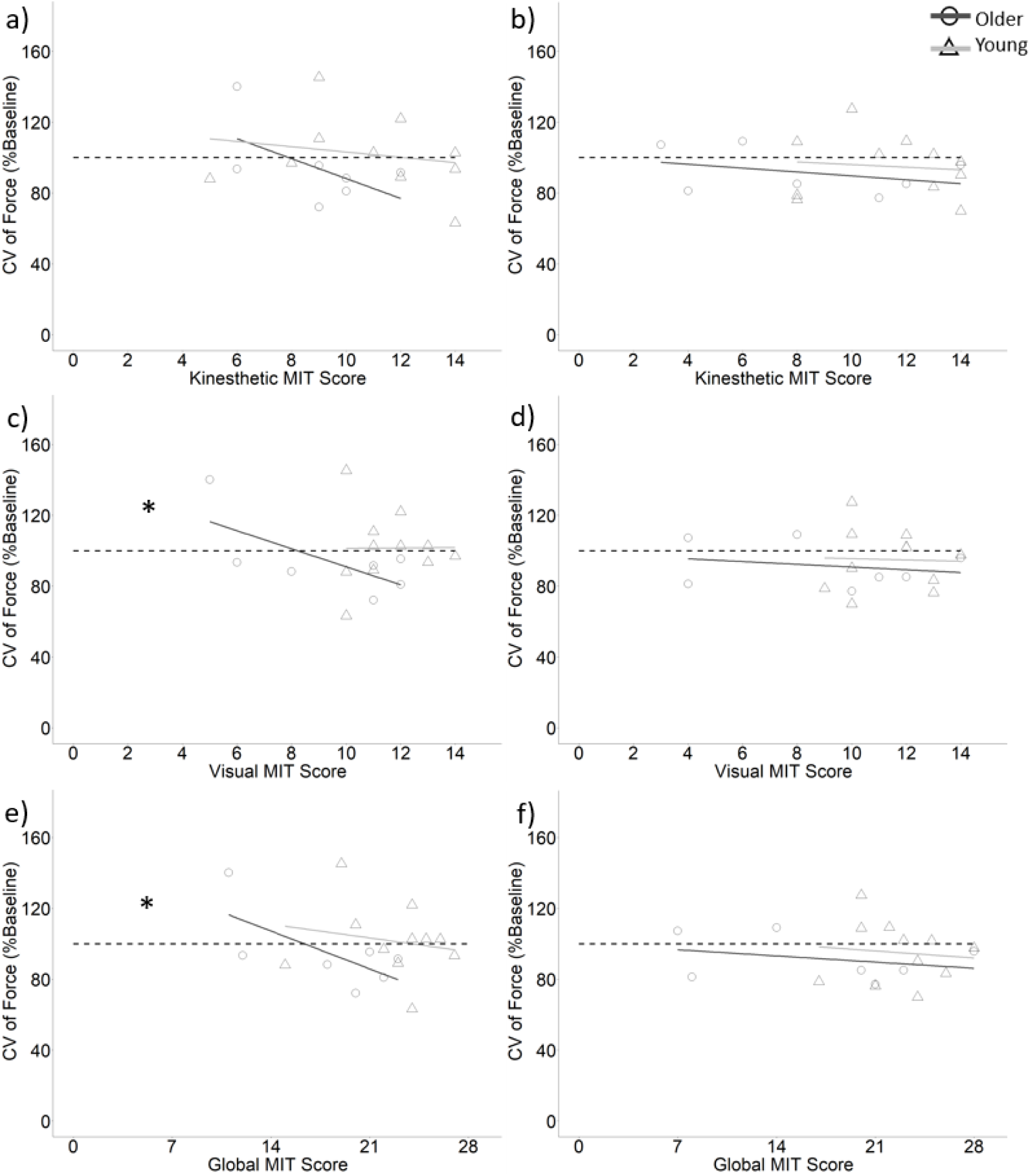
MIT scores from block 2 and CV of force from block 3 as a percentage of baseline for kinesthetic (a), visual (c), global (e) data. MIT scores from block 4 and CV of force from block 5 as a percentage of baseline for kinesthetic(b), visual (d), global (f) data. *, significant relationship for older females. MIT, motor imagery training; CV, coefficient of variation.

## Discussion

This study is the first to look at MIT priming in females, notably young and older females (i.e., above age 65), for a force-tracking task, and to highlight the effects of MIT priming specifically in understudied populations. One session of motor imagery training improved FS of a 10% isometric elbow flexion contraction in older and young females. In the older MIT group, FS improved after one block (i.e., a total of 10 minutes), and this improvement in FS was maintained following the second block of MIT. In contrast to the predicted increase of corticospinal excitability occurring concurrently with improved FS following MIT, corticospinal excitability decreased in the older MIT group after the second block of MIT. In the young MIT group, one block of MIT was insufficient to elicit a significant improvement in FS; however, following the second block of MIT, FS was improved compared to pre-training. In contrast to the older MIT group, no changes to corticospinal excitability occurred. This finding indicates that a total of 20 minutes of MIT improves FS in older and young females, with a concurrent decrease in corticospinal excitability only found in older females.

A novel aspect of this study is the evaluation of corticospinal excitability after MIT during a force-tracking task. Although prior studies have evaluated corticospinal activity during MIT (Kasai et al., 1997; Lebon et al., 2012; Lee et al., 2021; Mizuguchi et al., 2013), it is unknown whether corticospinal excitability is altered after MIT during a force-tracking task. In the older females, there was a significant reduction in MEP amplitude following 20 minutes of MIT during the tracking task. Kluger and colleagues (2012) found a decrease in MEP amplitude but not M-wave amplitude compared to baseline, at two minutes but not four minutes after kinesthetically and visually mentally rehearsing a maximal hand grip while participants (aged 30-59) were at rest. They suggest sustained central activation via motor imagery caused this decrease in MEP amplitude. Yet, future investigations are needed to explain why this is the case (Kluger et al., 2012). Therefore, the time between the given stimulus compared to when the MIT occurred could have influenced the level of corticospinal excitability during the tracking tasks. Following the MIT, participants rated their mental rehearsal and then performed the tracking task; this delay might have diminished any increased excitability induced by the MIT. The decline in MEP amplitude found only in older adults might also be consequential of MIT ability. Since kinesthetic and visual motor imagery were conducted together, it can only be speculated that reduced visual MIT ability in older adults compared to young may have influenced corticospinal modulation. As previously mentioned, greater motor imagery ability is associated with greater corticospinal excitability during MIT (Lebon et al., 2012). Even with high motor imagery ability in young females, the lack of change in corticospinal excitability might arise from the simplicity of the isometric task or the low submaximal intensity. Mizuguchi and colleagues (2013) reported greater corticospinal excitability in the biceps brachii and brachioradialis during MIT when imagining 60% MVC compared to 10% MVC. When imagining finger pinching compared to a finger tapping sequence, the more complex task of finger tapping induced higher corticospinal excitability than the simple task (Kuhtz-Buschbeck et al., 2003). Herein, young adults might require complex and challenging tasks to evoke changes in corticospinal excitability from MIT that will last acutely after training.

In this study, MIT priming likely contributed to an improvement in FS. After the first 10 minutes of MIT, there was a 5.4% improvement in CV of force relative to the baseline for the older training group, with no change for the Control group. This MIT priming effect was likely enhanced after an additional 10 minutes of MIT, as there was a further 3.0% improvement in CV of force from 10 minutes to 20 minutes of MIT. The compounding effect of two 10-minute blocks of MIT was an ∼8.4% relative improvement in CV of force.

In the review of neuroimaging studies, Munzert et al., (2009) reported that MIT activates the primary motor cortex akin to physical execution, although to a lesser extent. While, other meta-analytic work demonstrates inconsistent activation of the primary motor cortices during MIT (Hardwick et al. 2018; Hétu et al., 2013). The overlap of sensorimotor regions recruited during MIT and physical execution forms the basis for the simulation theory posited by Jeannerod (2001). Here, the mental rehearsal of a physical action (Jeannerod, 2001), creates a priming effect on force performance in young (Dos Anjos et al., 2022; reviewed in Stoykov & Madhavan, 2015). Di Rienzo and colleagues (2015) showed acute MIT priming, where isometric elbow flexor MVC force increased after training. MIT priming might be the result of increased neural excitability and/or a reduction in oscillations of common synaptic input (CSI) (i.e., shared input to the motor neuron pool). Age-related reductions of FS are associated with greater oscillations of CSI (Castronovo et al., 2018; Feeney et al., 2018; Pereira et al., 2019) and the amplitude in oscillations of CSI to motor neurons may also influence motor unit activation strategies to reach a target force level. Castronovo and colleagues (2015) observed that as contraction intensities increased from 20% to 50% to 75% of MVC, net excitatory input to motor neurons increased concurrently with the relative strength of CSI. Future studies should measure common synaptic input, to help identify the locus of neural change of the motor imagery priming on positively influencing force control. Considering all these findings, MIT priming has the potential to positively contribute to the mitigation of functional decline that occurs with aging, via improvement in force steadiness.

It is important to consider factors that may influence MIT priming, including MIT ability, and age. Here, we did not find evidence that kinesthetic, visual, and global MIT ability changed in older or young females between blocks. In the older MIT group, the global and visual motor imagery scores from the first 10 minutes of MIT were negatively related to the change in CV of force from pre-training; i.e., higher motor imagery scores related to greater improvement in CV of force and, therefore, a greater priming likely occurred. For the young MIT group, the CV of force was approximately 2% before and after 10 minutes of MIT, and this might suggest that, despite high MIT ability, there could be a ceiling effect in improving FS in young females.

FS improved with MIT in older females, despite the lower self-reported global and visual MIT ability relative to the young females. Malouin and colleagues (2010) suggested visual motor imagery is more affected by adult aging than kinesthetic motor imagery, with associated decreases in working memory. Our results support this hypothesis as self-reported visual ability scores were lower for older than young females, with no difference in kinesthetic ability scores. As visual and kinesthetic instructions were given to participants, the combined influence which is the global motor imagery ability when performing the isometric FS mental task likely influenced motor imagery priming. Future studies could look at visual motor imagery and kinesthetic motor imagery separately to understand the effects of each on improving FS in older adults. Even with reduced self-reported global motor imagery ability compared to the young, the older females demonstrated an ∼8.4% improvement in CV of force relative to the baseline after 20 minutes of MIT compared to the young in which a ∼5.0% improvement in CV of force relative to the baseline was observed. Ideally, to best understand age-related and sex-specific decline, future studies should include a large age-range cohort with both sexes. Consideration should also be given to measuring adults from ∼55 years, around when menopause occurs to establish rates of functional decline and the subsequent influence of MIT across age cohorts.

The current study contributed to knowledge on FS in females, specifically age-related declines in FS, which has received little consideration. Also, this study is the first to address MIT priming and FS in older females. The single session of MIT improved FS in older and young females, and older females decreased corticospinal excitability. Higher global and visual motor imagery ability during the mental rehearsal of submaximal steady contractions was related to improved FS for the older, but not young, MIT group. These findings indicate that older females benefit from MIT, and suggest this form of training might be useful to mitigate loss of FS with increased age. Therefore, the applicability of using MIT in real-world settings of older adults to improve functional ability is worthy of further investigation.

