## Supplementary Materials for "Motor Imagery Improves Force Control in Older and Young Females"

**Supplementary Appendix A. Motor Imagery Familiarization Script**

Motor imagery is the mental rehearsal of a physical task which means that you do not physically perform the task. Instead, you imagine you are performing the task by creating a picture of it in your head. For this study we want you to imagine yourself doing a physical task through your own eyes. Doing motor imagery can be difficult at first, but there are a few things that can help you get better at it. What you can do is try and relax by taking a couple of slow, deep breaths and letting yourself sink into the chair. As you are sitting, think about how the chair feels and the position of your body. Another thing you can do is to think about how it feels when you actually perform the task. How are you using your arm to pull up on the handle? How long does the task take? All of these sensations can be used to make the picture in your head more vivid. Right now, we want you to watch a short video so you can start to form a picture of yourself performing this studies task.

**Supplementary Appendix B. Motor Imagery Training Script**

Close your eyes. Imagine your surroundings through your own eyes. Imagine your arm placed in the chair rest and that you are holding onto the chair handle. Imagine the monitor screen in front of you and on the screen, there is the moving green force line. When you hear the word ‘up’ imagine the force line moving up as you pull up on the chair handle and feel your biceps muscle tighten. When you hear the word ‘hold’ imagine how much you have to exert yourself to make the green force line on the monitor screen stay at the target line. When you hear the word ‘rest’ keep your eyes closed, stop imagining and relax.
